# TCR2vec: a deep representation learning framework of T-cell receptor sequence and function

**DOI:** 10.1101/2023.03.31.535142

**Authors:** Yuepeng Jiang, Miaozhe Huo, Pingping Zhang, Yiping Zou, Shuai Cheng Li

**Affiliations:** Department of Computer Science, City University of Hong Kong, Hongkong, China

## Abstract

The T-cell receptor (TCR) repertoires are critical components of the adaptive immune system, and machine learning methods were proposed to analyze the TCR repertoire data. However, most methods work solely on the hypervariable CDR3 regions of TCRs, overlooking the information encoded in other domains. Representing full TCRs as informative vectors can be indispensable for developing reliable and effective machine learning models. We introduce TCR2vec, a deep representation learning framework with 12 layers of transformer blocks, to pave the way for downstream modelings of full TCRs. Together with masked language modeling (MLM), we propose a novel pretraining task named similarity preservation modeling (SPM) to capture the sequence similarities of TCRs. Through a multi-task pretraining procedure on MLM and SPM, TCR2vec learns a contextual understanding of TCRs within a similarity-preserved vector space. We first verify the effectiveness of TCR2vec in predicting TCR’s binding specificity and TCR clustering through comparison with three other embedding approaches. TCR2vec can be finetuned on small task-specific labeled data for enhanced performance, which outperforms state-of-the-art models by 2-25% in predicting TCR’s binding specificity. Next, we compare the performance of two versions of TCR2vec pretrained on full TCRs (TCR2vec) or CDR3s (CDR3vec) and demonstrate that TCR2vec consistently outperforms CDR3vec by 12-14%. Further analysis of attention maps reveals that residues outside CDR3 also make notable contributions to the recognition of antigens. TCR2vec is available at https://github.com/jiangdada1221/TCR2vec.

## Introduction

T cells are crucial to immune protection through their ability to recognize diverse pathogens specifically and to provide enhanced protection against reinfection. The receptors (TCRs) on their surface need to recognize an immunogenic peptide presented in the context of major histocompatibility complex molecules (MHC) to activate T cells in an immune response. The generation of TCRs is accomplished via the process of somatic recombination of V, D (*β* chain only), and J gene segments, which can produce an extremely high TCR diversity of 10^15^ −10^20^ variants in an individual ([1]). Each TCR is composed of an *α* chain and a *β* chain, both of which possess three complementary-determining regions (CDRs). These hypervariable regions largely determine the antigen specificity of the TCR.

Machine learning (ML) models have been used to analyze TCR sequences (repertoires) and achieved great success in various applications, such as predicting their recognition specificity ([2, 3]), deciphering their sequence patterns ([4, 5]), and inferring the thymic selection pressures during T cell maturation ([6]). Relying on the power of ML models to learn the patterns of the data, we can now predict the outcomes of complex biological processes without prior knowledge of their underlying mechanisms. To develop a ML model for TCRs, one must embed the TCR sequence into a numerical representation compatible with the arithmetic calculations applied in ML models. A key factor in the performance of ML models is the quality of the input numerical embeddings. Powerful prediction models can produce poor results with less informative embeddings ([7]). Thus, high-quality embeddings of TCRs are in demand and can facilitate the development of ML models for TCRs.

In TCRs, CDR1 and CDR2 typically contact the conserved *α*-helices of MHC ([8, 9]), while CDR3 is the main determinant responsible for the recognition of the processed antigen ([10]). So far, most work focuses on analyzing the CDR3 of the TCR-*β* chain. However, the exclusion of CDR1 and CDR2 would completely ignore their roles, though not as important as CDR3, in antigen recognition and might tamp down the reliability of conclusions drawn solely by CDR3. In addition, the constant regions can also affect the peptide binding ([11, 12]). Thus, to fully capture and unravel the function of each TCR, all regions should be included for reliable conclusions to be drawn.

A key hurdle impeding the development of ML models for full-length TCRs is the lack of a suitable representation learning framework that can transform them into informative vector representations that capture the function of each TCR within the context of the large TCR universe. Some of the previous forms of CDR3 representation learning focused on calculating the physicochemical properties of CDR3 sequences ([13]) or extracting the hidden nodes from autoencoder models ([4, 5]). Though these methods have been successfully used for binding specificity prediction and repertoire classification, directly applying them to full-length TCR sequences remains challenging as they have to extract meaningful information from a larger and more complicated context (roughly ∼120 amino acids in a TCR with ∼15 amino acids inside CDR3 region). Some models attempted to somehow incorporate knowledge from other domains by taking as input the CDR3 sequences as well as their corresponding V and J genes ([4, 6]). However, they embed the genes via the one-hot representation, which might lose much information encoded in their sequences.

Recently, deep learning models have demonstrated promising results in deciphering the hidden patterns of amino acid sequences that correspond to their function and structure ([14, 15]). Neural networks, especially transformers, are adapted from the text domain to the protein domain by treating amino acids as “words” and protein sequences as “sentences”. The self-supervised pretraining on massive data largely contributes to their success. A common strategy of pretraining is the masked language modeling (MLM), where a model learns the language structure by predicting missing parts of the protein sequence conditioned on the peripheral contexts. General protein language models (LMs), such as ProteinBERT ([16]) and ESM ([14]), can then generate a distributed embedding of each amino acid in the context of the protein sequence, acting as a “warm” starting point for downstream tasks including protein engineering and protein structure prediction ([17, 18]).

The motivation behind these large-scale LMs is that functions are encoded in the sequence through a complex manner. Traditional hand-crafted embeddings may miss the latent sequence patterns in the data that are not easy to observe and distinguish directly. However, by feeding abundant unlabeled data to LMs, they can automatically and efficiently learn the sequence language. The transformer blocks with the multi-head attention mechanism provide LMs the capacity to understand the relationship between sequential elements that are far from each other and thus produce informative embeddings for input sequences. Note that in most applications of ML models, domain-specific models can bring improvement to general models. For example, models for specific protein families can outperform general protein models ([19, 20]). Likewise, B-cell receptor (BCR) specific LMs can offer superior performance to general protein LMs ([21]). Therefore, to pave the way for the usage of full-length TCRs, a TCR-specific LM that can capture the nuances of TCRs is of great significance.

Though MLM has demonstrated effectiveness across different LMs in learning contextual representations, it is not sufficient to extract the functional information encoded in the similarities between TCRs. Despite the complex and unique nature of TCRs, TCRs that recognize the same antigen frequently have highly similar sequences or binding patterns (motifs) ([22–25]). Thus, embedding approaches that can reflect sequence similarity in the vector space may facilitate function-related computational tasks. For example, GIANA ([26]) is a TCR clustering framework based on embeddings of CDR3 sequences with the characteristic of sharing high linear correlations between pairwise Euclidean distances and sequence similarities. However, this correlation only holds for pairs of the same length. A proper method is required to preserve the pairwise similarities between amino acid sequences in embedding space.

In this work, we present TCR2vec to fill the gap of the TCR-specific language model. TCR2vec is a deep representation learning framework specific for T-cell receptors based on a 12-layer transformer model that is pretrained on 1 million human TCR sequences. Meanwhile, we propose a new pretraining task, similarity preservation modeling (SPM) that aims to preserve the pairwise sequence similarities of TCRs in embedding space. By jointly optimizing MLM and SPM with multi-task learning, TCR2vec can generate embeddings for TCRs that simultaneously encode contextual and functional information. We showcase the usefulness and robustness of TCR2vec in two important downstream tasks: prediction of antigen specificity (supervised task) and TCR clustering (unsupervised task). Therefore, TCR2vec could serve as a basis for TCR-specific representation learning, a key step in computational immunology applications.

## Methods

### Reconstruction and annotation of full TCR sequences

The first step in data curation is the reconstruction of full-length TCRs based on the CDR3 sequences as well as their corresponding V/J genes. We followed the reconstruction process in [27]: the end of the V gene segment (last five amino acids) was aligned to the CDR3 sequence and then the V segment was merged with the CDR3 sequence based on the alignment results. Similarly, the CDR3 sequence was first aligned to the J gene segment and then merged with the J segment. Based on the two merging results, we can assemble these segments to reconstruct the full-length TCR sequence. The full-length TCRs were annotated by ANARCI ([28]) using the IMGT numbering scheme ([29]) to identify each CDR region.

### Data

To construct comprehensive TCR sequence data for the pretraining stage, we combined TCR repertoires sampled from a large cohort of 743 individuals collected in [30]. We performed some filtering steps for this dataset, including the removal of sequences with unresolved V or J genes and the drop of sequences with ambiguous amino acids (*B, J, O, U, X*). Sequences that failed in the full-length TCR reconstruction were also excluded. Finally, we pooled the unique TCRs from each repertoire and constructed a large TCR pool of ∼90M TCRs. We randomly sampled 1 million TCRs from this pool for pretraining TCR2vec and refer to this subset as *Emerson data*. We constrained ourselves to TCR-*β* chain in this work, given that the paired data is still scant.

To evaluate the performance of TCR2vec in the tasks of predicting the binding specificity and TCR clustering, we combined the data recording the links of TCRs to their epitope targets collected in VDJdb ([31]), McPAS-TCR ([32]), and IEDB ([33]). The same filterings for *Emerson data* were first performed to the combined epitope-specific TCR dataset. Then, only human TCR sequences were selected; duplicates and cross-reactive cases were removed; only MHC class I entries with linear epitope sequences whose lengths lie between 7-15 amino acids were kept. We also removed rare epitopes with less than 10 recorded links to TCRs. At last, we were left with a dataset of 100,112 pairs of TCRs with binding targets of 369 epitopes. Given that these TCR-epitope datasets contain only positive samples, we need to sample negative samples, as both positive and negative samples are required to evaluate the model performance in the supervised task. The negative pairs were generated based on the *Unified Epitope* strategy suggested in [2] via shuffling the positive pairs: for each TCR in the positive data, we sampled an epitope from the positive data as its negative target according to the frequency distribution of epitopes while excluding its true epitope targets. Finally, we adopted the standard 5-fold cross-validation process to evaluate the performance in the supervised task.

### The proposed TCR2vec framework

#### Model architecture

TCR2vec is a TCR-specific Bidirectional Encoder Representation from Transformers (BERT; [34]). Specifically, we adopted the modified BERT model (TAPE-Transformer) for proteins introduced in [15]. We set the model parameters the same as the *BERT-base*, except that we used a smaller hidden size of 120 for computation efficiency since we did not observe notable improvement for the original hidden size of 768. There are 12 hidden layers with an intermediate feed-forward size of 3072. The number of the self-attention heads is 12. Using this BERT model, a batch of input TCRs is converted into a 3D tensor **X**_*out*_ ∈ ℝ^*B×max*(*L*)*×d*^, where *B* is the batch size, *max*(*L*) is the maximum sequence length in the current batch, and *d* is the hidden size which is 120 here. Following [21] and [14], we averaged the output tensor over the length dimension while excluding the padding tokens to derive the fixed-sized embeddings **X**_*emb*_ ∈ ℝ^*B×d*^ for the batch of TCRs.

#### Similarity preservation modelling

To explicitly enforce embeddings to retain the pairwise similarity in sequence space, we designed a new pretraining task called similarity preservation modeling (SPM). The general workflow of SPM is illustrated in Fig. 1. For a batch of *N* TCRs *T*_1_, …, *T*_*N*_ along with their embeddings 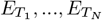, we first defined the following two vectors *V*_*seq*_ and *V*_*emb*_ as:

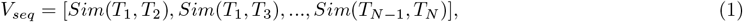

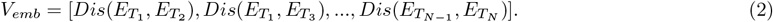

**Figure 1:**
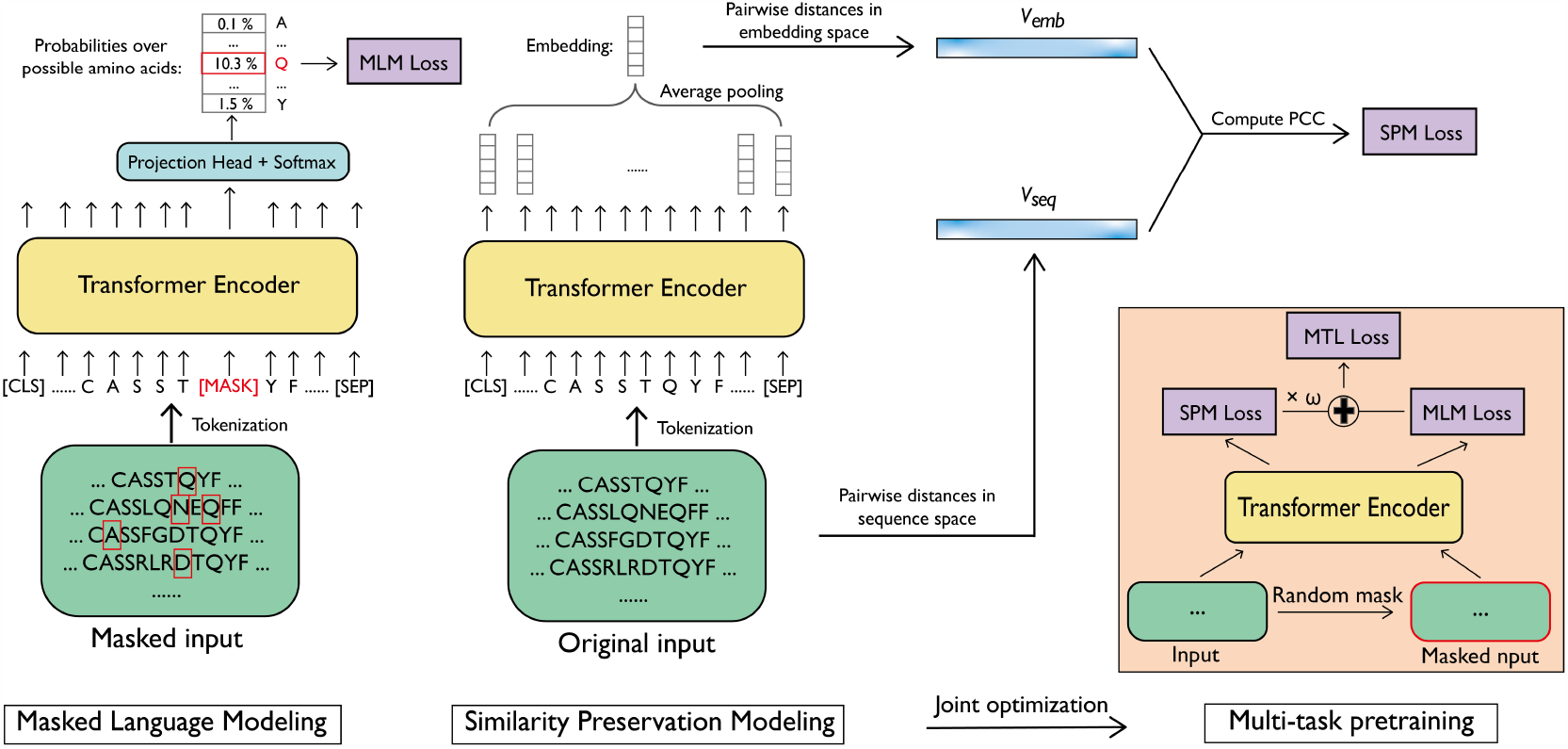
The pretraining workflow of TCR2vec. In MLM, the input TCR sequences are corrupted based on the random masking schema and the model learns to predict the correct amino acid for the masked positions (left panel). In SPM, the model is trained to retain the sequence similarity in the embedding space. i.e., pairs of TCRs with higher sequence similarities possess smaller distances in embedding space (middle panel). TCRvec adopts a multi-task pretraining procedure that jointly optimizes MLM and SPM to learn the contextual understanding of TCRs in a similarity-preserving space (right panel).

**Figure 2:**
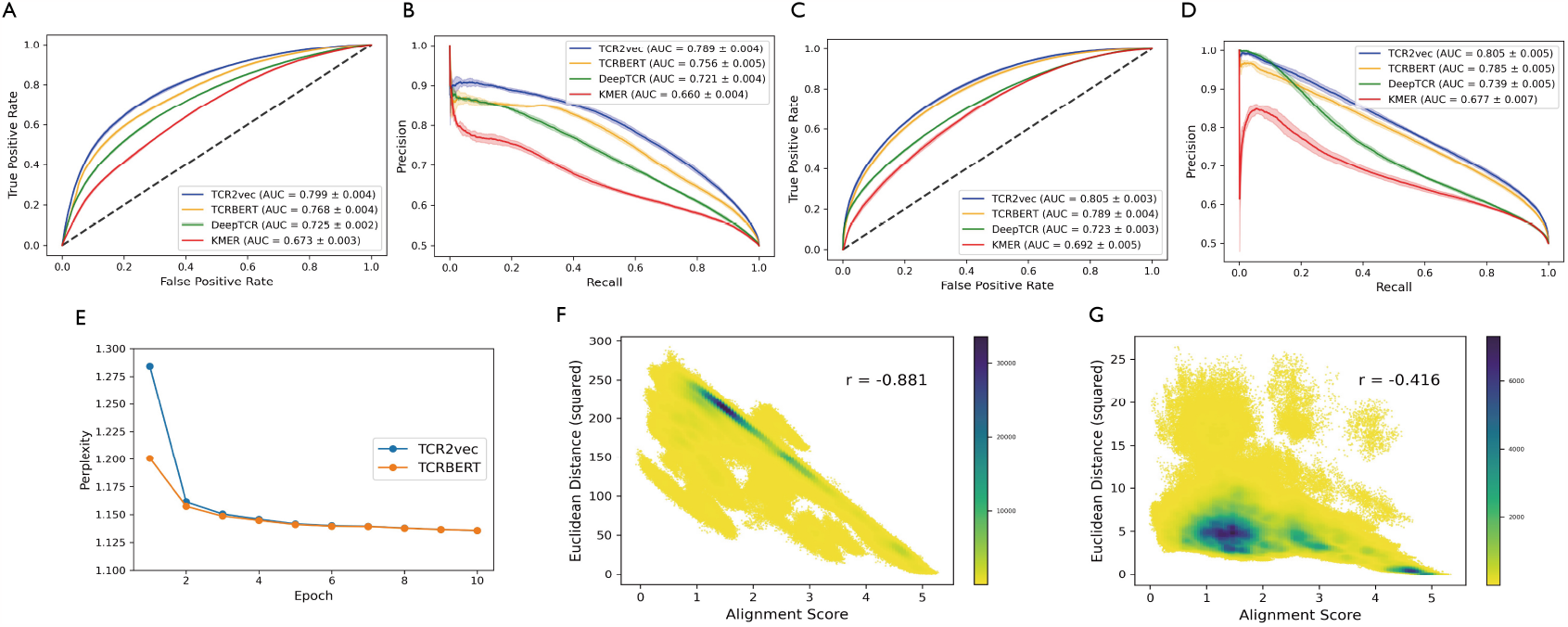
The prediction and pretraining performance of TCR2vec. (A) and (C) The ROC curves with the respective classification heads SVM and MLP. (B) and (D) The PRC curves with the respective classification heads SVM and MLP. These classification results are based on the *EigDecom* method for embedding epitopes. (E) The perplexity of the validation set during the pretraining process. (F) and (G) The correlation between the sequence similarities (measured by alignment scores) and the distances in embedding space with (F) or without (G) the SPM in the pretraining procedure. These values are computed on the validation TCRs.

*V*_*seq*_ denotes the vector of the pairwise similarities between the batch of TCR sequences measured by the normalized Needleman-Wunsch alignment scores using the BLOSUM62 substitution matrix ([35]); *V*_*emb*_ denotes the vector of the pairwise squared Euclidean distances of those *N* TCRs in the embedding space. Then, in the task of SPM, TCR2vec is optimized to lower the correlation between *V*_*seq*_ and *V*_*emb*_, which is equivalent to maximizing the negative correlation between *V*_*seq*_ and *V*_*emb*_. The loss function can be formulated as follows:

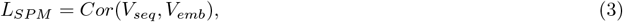

where *Cor*(*V*_1_, *V*_2_) computes the linear correlation between two vectors *V*_1_ and *V*_2_; we used Pearson correlation coefficient (PCC) to quantify such correlation. The reason for minimizing the *L*_*SP M*_ is that the PCC values are negative since larger alignment scores implicate higher sequence similarities, which correspond to smaller distances in the embedding space. This pretraining objective guides the TCR2vec to embed TCRs into a similarity-preserved vector space, inside which similar TCRs have small Euclidean distances and meanwhile, divergent TCRs have large Euclidean distances.

#### Multi-task pretraining procedure

The masked language modeling is commonly used to pretrain transformer-based models, enabling the models to learn a good contextual understanding of an entire sequence (Fig. 1). In addition to SPM, we also applied MLM for pretraining TCR2vec to learn the complementary contextual information of TCR sequences. To fully exploit the learning ability of MLM and SPM, we proposed a multi-task pretraining procedure to jointly optimize the objectives of MLM and SPM. As shown in Fig. 1, the 3D output tensor **X**_*out*_ was used for MLM, and the embedding tensor **X**_*emb*_ was applied for SPM. Then, the final loss function for TCR2vec is formulated as a multi-task objective:

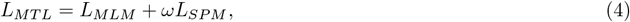

where *ω* represents the task weight of SPM.

### Evaluation

The performance of TCR2vec was tested in two downstream tasks that are closely related to the function of TCRs. The first task is the generic TCR-epitope binding prediction task, where a model predicts whether or not a pair of TCR and epitope can interact. Previous works have attempted to solve this challenging task by unlocking the binding pattern between TCRs and epitopes, such as ImRex ([13]), TITAN ([27]), and TEINet ([2]). A common model design for this generic classification task is the two-tower architecture: two separate encoders produce respective embeddings of TCRs and epitopes, which are concatenated and input into a classifier to output the binding scores. Following this concept, we adapted two embedding methods introduced in [3] and [36] to encode the epitope sequences into vectors with the same vector size as TCR2vec (Supplementary Text S1). We refer to these two epitope-embedding methods as *EigDecom* and *AtchleyAE*. Then we concatenated the embeddings of TCRs and epitopes and applied the Support Vector Machine (SVM) ([37]) or the Multi-layer Perceptron (MLP) classifier for prediction. The classification performance is quantified by the area under a receiver operating characteristic curve (AUROC) and the area under the precision-recall curve (AUPRC). Higher AUROC or AUPRC values demonstrate better classification performance, indicating that the embeddings are more informative with respect to the functional properties of TCRs.

In addition to the supervised classification task, we further evaluated the performance of embeddings in an unsupervised task of clustering TCRs. Ideally, TCRs specific to the same epitope should be included in the same cluster. Despite the huge success achieved by the *k*-mer based clustering methods such as GLIPH1 ([23]) and GLIPH2 ([38]), recent works in TCR clustering such as GIANA ([26]) and ClusTCR ([39]) have shifted their focuses towards embedding-based clustering. Another branch of models clusters TCRs according to their pairwise distances computed by their amino acid compositions, such as TCRdist ([22, 40]). A major shortage of TCRdist is the high time cost, as it has a quadratic time complexity for performing the pairwise comparison. However, this can be alleviated by the transition from sequence space to embedding space. The pairwise comparison regarding the Euclidean distances in embedding space can be computed via vectorization, making it computationally efficient and scalable to large datasets. To summarize, we evaluated the utility of TCR2vec in TCR clustering from two facets: (1) we first used the distance-based clustering algorithm (e.g., TCRdist) to cluster TCRs, with pairwise distances computed by the Euclidean distance in embedding space; (2) we adopted the embedding-based clustering framework GIANA and replaced its inside embedding method with TCR2vec. The performance is quantified by the clustering purity under different retention values. Under specific retention values, higher purity values indicate better clustering performance. They are defined as previously described in [41]. The retention is formulated as the total number of TCRs *T*_*i*_ that participate in forming any cluster *c* divided by the data size *N* :

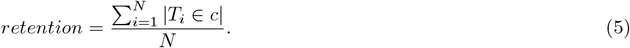

Purity is defined as the proportion of sequences within any cluster possessing the same epitope specificity. For each cluster *c*, the most common epitope *e*_*c*_ is considered the specificity of this cluster. We count the number of TCRs inside each cluster *c* specific for *e*_*c*_, sum the values, and divide them by the total number of sequences (*N*_*c*_) in any cluster:

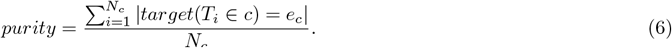

### Implementation and Training details

For details of MLM, 15% amino acids within the TCRs are randomly selected for masking. In 80% of the cases, the masked amino acids are replaced by a special token “[MASK]”; in 10% cases the masked tokens are replaced by random amino acids that are different from the original ones; in the remaining 10% cases, we kept the original amino acids. For the training of TCR2vec, we mainly followed the settings defined in [15]: TCR2vec was trained for 10 epochs with a batch size of 64 and a dropout rate of 0.1 on *Emerson data*. The optimizer used is Adam ([42]) with a learning rate of 10^*−*4^ and a linear warm up schedule. We empirically set the task weight of SPM (*ω*) to 10, as it performed well on the downstream classification task. The NeedlemanWunsch alignment scores were computed using the Parasail library ([43]) with an open-gap penalty of 6 and an extend-gap penalty of −1. We normalized the pairwise alignment score by dividing the maximum length of the two input sequences. For the classification heads in the prediction task, the SVM with the RBF kernel and MLP were implemented by the sklearn library ([44]). We set the regularization parameter *C* in SVM to 10; the MLP has 2 hidden layers with respective 256 and 128 neurons and was trained for 50 epochs with a batch size of 128. 15% of the training data was set aside as internal validation set to perform early-stopping. Other parameters not mentioned were set to default values.

## Results

### TCR2vec outperforms other embedding approaches in the classification task

We first compared the quality of predictions using the SVM and MLP with different embeddings. We selected three baseline embedding approaches for comparison: a *k*-mer based method from [45] (we refer to it as “KMER” in the following text); the variational autoencoder (VAE) model DeepTCR from [4] that encodes the gene information as one-hot vectors; TCR2vec without the SPM pretraining, which serves as a regular BERT-like model (referred to as “TCRBERT”). For a fair comparison, their embedding sizes are restricted the same as TCR2vec. Detailed descriptions of the baseline models are included in Supplementary Text S2. Based on the comparison to these models, we aimed to show that: (1) TCR2vec can outperform the *k*-mer approach that was commonly used before the emergence of large-scale language models; (2) instead of encoding the V and J genes as one-hot vectors by their names, modeling the amino acids within gene segments are more useful; (3) SPM pretraining is helpful for extracting the functional information encoded in TCR sequences.

Figure 1A-1D show the classification performance of each model using the *EigDecom* for encoding epitope sequences. We observed that TCR2vec achieved respective AUROCs of 0.799 ± 0.004, and 0.805 ± 0.003 with SVM and MLP classifier, outperforming its competitors. Besides, for the metric of AUPRC, TCR2vec again demonstrates higher values than baseline methods (0.789 ± 0.004 and 0.805 ± 0.005 with SVM and MLP). We noticed that the KMER method performs the worst (0.673 ± 0.003 and 0.692 ± 0.005 AUROCs; 0.660 ± 0.004 and 0.677 ± 0.007 AUPRCs). The low performance of KMER may be attributed to the lack of contextual understanding of TCRs as sub-units (*k*-mers) of the amino acid sequences are considered independent. Thus, their covariances with the remainder of the TCR sequence are overlooked. DeepTCR performs better than KMER (0.725 ± 0.002 and 0.723 ± 0.003 AUROCs; 0.721 ± 0.004 and 0.739 ± 0.005 AUPRCs) since the middle layer of VAE can serve as a summative and compact representation for the TCR. However, it is largely surpassed by TCR2vec, indicating that the one-hot encoding of genes is less informative than directly handling their corresponding amino acids. Finally, TCR2vec performs better than TCRBERT (0.768 ± 0.004 and 0.789 ± 0.004 AUROCs; 0.756 ± 0.005 and 0.785 ± 0.005 AUPRCs), suggesting the usefulness of SPM in learning more informative embeddings. Besides, using MLP improves upon SVM for TCRBERT (from an average AUROC of 0.768 to 0.789), demonstrating the necessity of using a more complex model (MLP) to extract the function information encoded in embeddings. In contrast, TCR2vec obtains similar prediction results when applying either SVM or MLP, indicating that its embeddings are informative such that a simple classification head is enough to achieve high accuracy. Similar results are observed when using the *AtchleyAE* for embedding epitopes (Supplementary Figure S1).

To resolve the concern that SPM might negatively impact the MLM during the multi-task pretraining process, we show the training curve of TCR2vec and TCRBERT in Fig. 1E. On the one hand, we found that with SPM, the perplexity (the metric for evaluating MLM) can ultimately decrease to the same level of using only MLM. On the other hand, TCR2vec can achieve a high level of correlation between the sequence similarities and distances in embedding space compared to TCRBERT (Fig. 1F and 1G). Thus, the multi-task learning process enables TCR2vec to preserve the sequence similarity while not sacrificing its contextual understanding of TCRs.

### Comparison to state-of-the-art TCR-epitope binding prediction models

TCR2vec was further evaluated by comparing its performance in predicting TCR-epitope binding specificity with state-of-the-art models. We finetuned TCR2vec on this task with a two-layer MLP for prediction to further exploit the capacity of TCR2vec (Supplementary Text S3). We selected ImRex ([13]), TEINet ([2]), and TITAN ([27]) for comparison. ImRex and TEINet both take the CDR3 sequences as input, which are embedded by either the physicochemical properties (ImRex) or an autoencoder model (TEINet). Similar to TCR2vec, TITAN utilizes the full-length TCR sequences which are encoded into hand-crafted embeddings using the BLOSUM62 matrix.

Figure 3 shows the results of TCR2vec and the comparative models. We first observed that TCR2vec largely outperforms ImRex and TEINet with respective enhancements of 25.4% and 11.9% regarding the AUROC. This result consolidates the promise of using full-length TCR sequences to capture the full complexity encoded in TCRs. Moreover, when using a less efficient embedding method in TITAN, the performance is lower than TCR2vec (AUROC = 0.840 ± 0.003 of TCR2vec compared to AUROC = 0.824 ± 0.004 of TITAN). Note that TITAN adopts a complex convolutional neural network (CNN) model with the attention mechanism to process the input embeddings. This further manifests the significance of the representation learning for TCRs, since even with an advanced deep-learning model, less informative embeddings would produce less accurate results.

**Figure 3:**
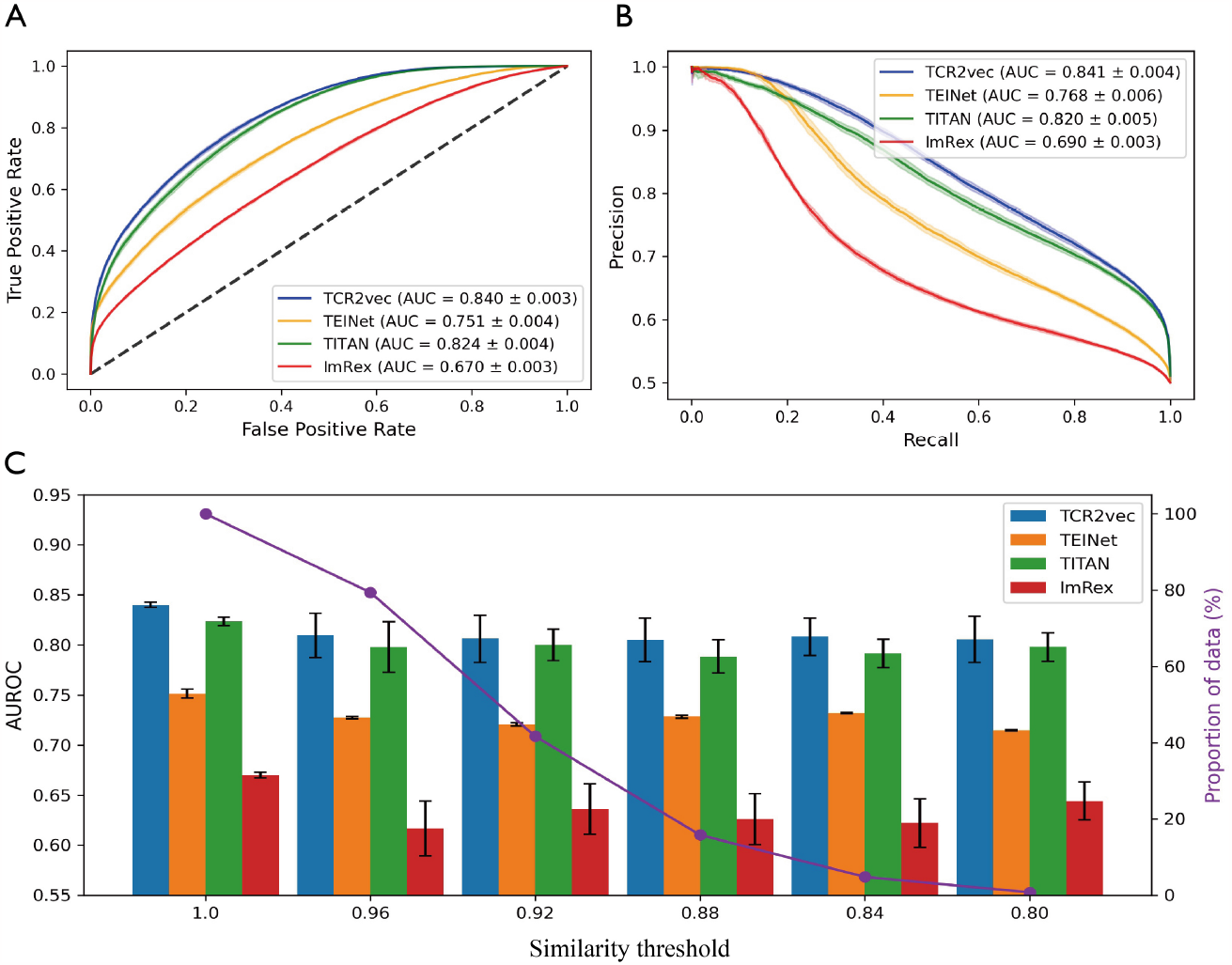
The performance comparison between TCR2vec and SOTA methods. (A) The ROC curves of each method. (B) The PRC curves of each method. (C) The AUROC for each model according to different similarity thresholds for filtering the test set. The purple curve shows the proportion of the filtered data to the original data.

At last, we investigated whether the superiority of TCR2vec retains for TCRs in the test set with decreased similarity to TCRs in the training set. This was done by filtering out pairs in the test set with TCRs exceeding specific similarity scores with any of TCRs in the training set (Supplementary Text S4). The classification performances of each model under different similarity thresholds are shown in Fig 3C. TCR2vec again performs robustly for different thresholds, indicating its good generalization capability.

### TCR2vec can facilitate TCR clustering

We next assessed TCR2vec and other embedding methods in the task of TCR clustering. Based on TCRdist, we first utilized the pairwise distances in embedding space for clustering to directly infer the quality of embeddings. We show the values of purity under different retention values in Fig. 4A. Firstly, all embedding methods except KMER demonstrate improvements over TCRdist which calculates the distances based on the amino acid sequences and the corresponding V genes ([22]). Thus, clustering TCRs based on the pairwise distance in vector space can not only boost the performance but reduce execution time via vectorization. Among the embedding approaches included, TCR2vec achieves the best clustering performance, with the purity-retention curve overlying other methods. Both TCR2vec and TCRBERT outperform DeepTCR, indicating the information enrichment from the use of full amino acid sequence and the superiority of LM-based models. Besides, TCR2vec again surpasses TCRBERT, illustrating the effectiveness of embeddings that are able to preserve sequence similarity.

**Figure 4:**
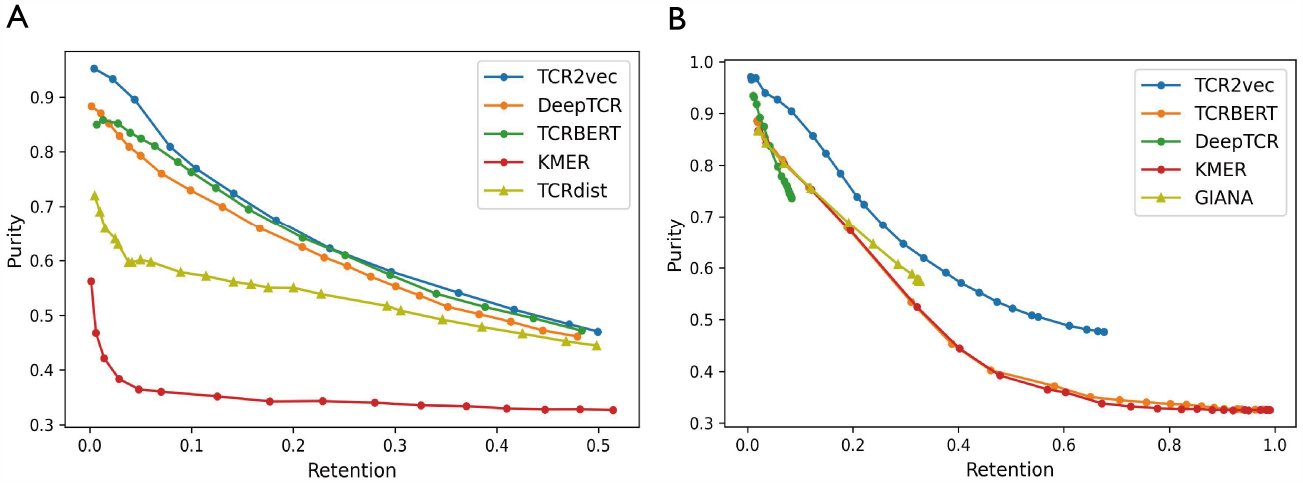
The performance in the task of TCR clustering. (A) The purity-retention curves of each embedding method for the distance-based TCR clustering method. (B) The purity-retention curves for the two-step TCR clustering method GIANA with its embeddings substituted by each embedding method.

To further compare each embedding method in TCR clustering, we utilized the embedding-based clustering framework GIANA ([26]) with its original embedding substituted by TCR2vec and other embedding methods. GIANA is a representative of the two-step TCR clustering framework in which TCRs are first grouped into “pre-clusters” using conventional clustering algorithms such as *K*-means on their embeddings, and then these pre-clusters are further refined in the next step. Thus, the quality of the embeddings influences those pre-clusters directly, which further impacts the cluster refinement afterward. We changed the critical parameter “S” in GIANA to control the level of retention and restricted GIANA to only use the sequence information. Note that for each embedding method, the minimum value of “S” was set low enough that no further increase in retention was observed. We show the clustering results with the GIANA framework in Fig. 4B. The superiority of TCR2vec is further underpinned, as it outperforms other embedding approaches by a large margin. Compared to DeepTCR and the original GIANA, TCR2vec gives a more “broad” clustering performance with enhanced purities, as DeepTCR and GIANA limit themselves to a small range of retention values. Note that TCRBERT performs as poorly as KMER. Though they possess a broader range of retentions compared to TCR2vec, they both demonstrate significantly decreased purity values. This observation showcases the indispensability of SPM in learning informative embeddings, which contributes to the robustness of TCR2vec across different evaluation settings to a large extent.

### Limited functional information is encoded in the CDR3 region of TCR

We further explored the functional information gained when using the full-length TCR compared to only the CDR3 region. For a more direct comparison, we re-trained TCR2vec on CDR3 sequences and obtained a CDR3-specific TCR2vec model, which we call “CDR3vec”. Figure 5A - 5C show the performance comparison between CDR3vec and TCR2vec in the prediction and clustering tasks. As expected, TCR2vec achieves better performance than CDR3vec in both tasks, implying that the CDR3 region only partly encodes the function of the TCR.

**Figure 5:**
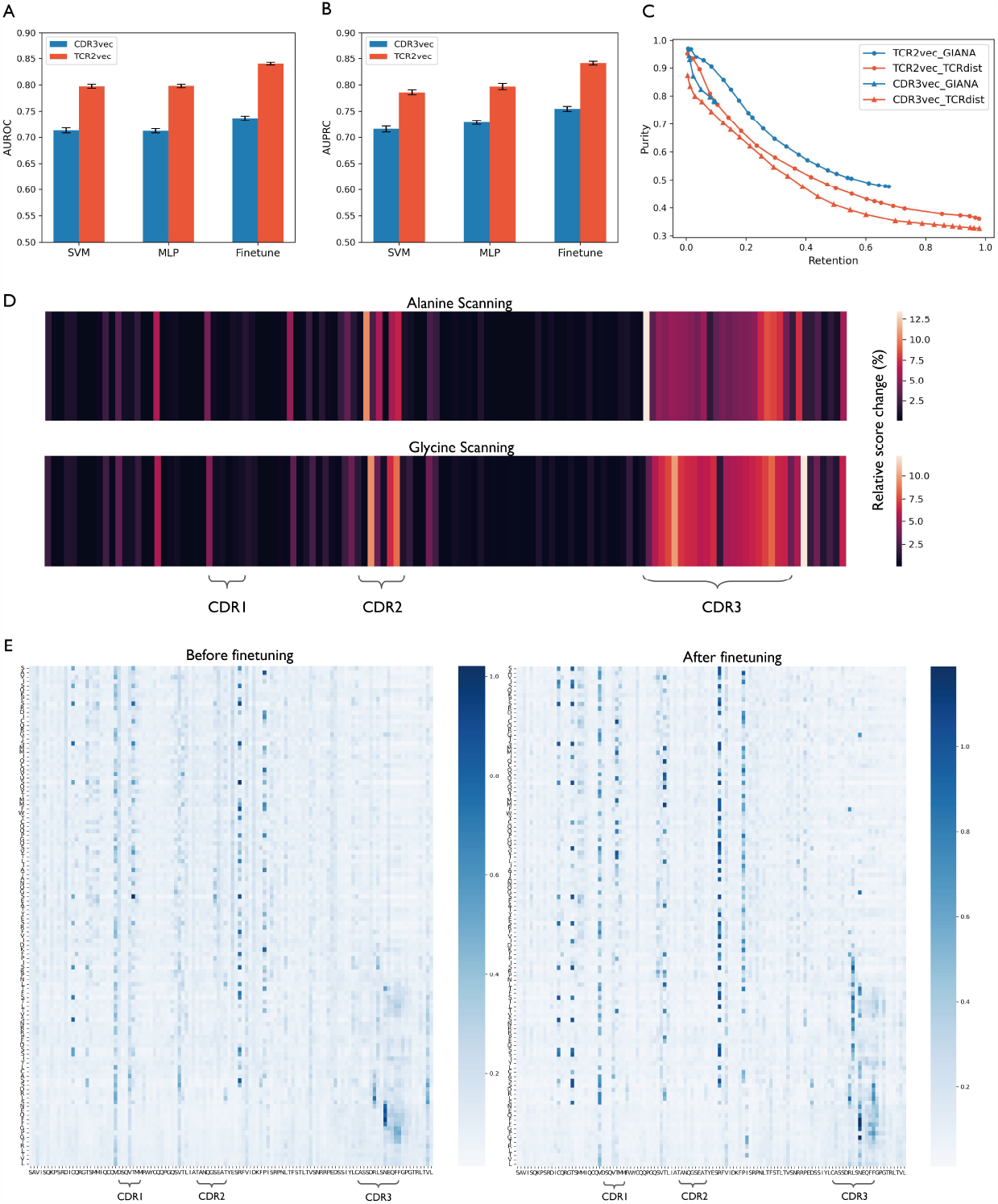
Comparison between the TCR2vec models pretrained on CDR3 sequences (CDR3vec) and full length TCRs. (A) The AUROC values for each model using the SVM or MLP classification heads. We also include the results of finetuning the CDR3vec or TCR2vec. (B) The AUPRC values for each model under different settings. (C) The clustering results of CDR3vec and TCR2vec using the distance-based clustering framework (TCRdist) or the two-step clustering framework (GIANA). (D) The per-residue relative score change when performing the alanine (top panel) or glycine (bottom panel) scanning technique. Details of the scanning procedure are included in Supplementary Text S5. (E) The self-attention heatmap from the last layer of TCR2vec before and after finetuning. We summed the attention map of each head. The direction of attention is from positions in the rows toward positions in the columns. Darker shades of blue indicate stronger attention.

To examine the contributions of each domain within TCRs to antigen recognition, we performed the alanine/glycine scanning technique used in biophysics studies ([46]) and previous works ([2, 5, 36]) to quantify the contributions (Supplementary Text S5). Alanine/glycine scanning is a simulated mutation analysis designed for obtaining structural evidence of TCR residues whose mutations can lead to significant changes in the predicted binding strength between the TCR and epitope. By virtue of the scanning approach with the trained prediction model, we observed that the CDR3 domain is the most prominent region relating to the peptide binding, as the majority of residues can lead to a large change in the prediction score when being substituted by alanine or glycine (Fig. 5D). Additionally, some important residues inside the CDR1 and CDR2 regions are detected, indicating that they also play an important role in determining the TCR’s function. These findings are in accordance with previous studies ([3, 22]) that CDR1-3 regions synergistically drive the antigen recognition of TCRs with the CDR3 region playing a more crucial role. Moreover, we also observed some prominent residues with large alterations of prediction scores in conserved regions of the TCR (e.g., residues near the CDR2 region). Thus, the contributions from constant domains should not be overlooked.

We further interpreted the per-residue contribution of the TCR sequence using the self-attention scores. These scores provide clues on deciphering the importance of the inner interactions in the final contextual embedding for each amino acid within the TCR sequence. We extracted the attention maps from the last layer of TCR2vec before and after finetuning on the classification task and summed them across each attention head to obtain a general attention map. We show an example in Fig. 5E. Before finetuning, the attention map is sparse but with some notable concentrations on residues around CDR1, the right side of CDR2, and within CDR3. After finetuning, the attentions of these regions are further amplified, indicating that they are more decisive for the TCR-epitope binding. Besides, we noticed that after finetuning, some residues inside CDR3 begin to gain large attention scores for amino acids preceding it. This finding manifests that the interactions between CDR3 and some preceding residues also contribute to the determination of the TCR’s specificity. Thus, using CDR3 sequence solely would not only loss useful information from other parts, but also overlook such inner interactions. More examples of the attention maps are available in Supplementary Figure S2.

## 1 Discussion

Recent advancements in high-throughput T-cell receptor sequencing technologies have enabled the generation of an increasing amount of TCR repertoires. The massive unlabeled data provides an access to learning the language of TCRs encoded within the amino acid sequence. In addition, owing to the broad clinical applications of TCRs, investigating the TCR sequence data through computational modeling has been an active research field and has illustrated meaningful biological insights ([22, 23, 36]). Machine learning models have stood out and served as powerful tools to unlock the complex pattern governing the recognition ability of TCR. The development of a representation learning framework for TCRs that can extract the latent functional information in sequence space and encodes it into embedding space is critical to the success of ML models.

Previous studies mostly concentrated on the CDR3 region of the TCR, as it is more likely to be in close contact with peptides. Although these CDR3-based works have achieved great success in bioinformatics tasks such as predicting the binding of TCR and epitope ([2, 36]), clustering TCR into groups with the same specificity ([23, 26]), inferring the thymic selection pressure ([6]), and cancer diagnosis ([47]), their performance could be further improved by including information from other parts of the TCR. Some works have shown that all CDRs contribute to the epitope recognition ([3, 22]). Additionally, the constant domains are also found to influence the peptide binding ([11, 12]). Our experimental results of alanine/glycine scanning and analysis of the attention maps also confirm these findings. Thus, to fully capture the complexity of TCRs and draw more reliable and comprehensive conclusions, we advocate the usage of full-length TCR instead of only the CDR3 region.

To establish the usage of full-length TCR, we present TCR2vec in this work. TCR2vec is a BERT-based representation learning framework that deciphers the sequence pattern of TCRs and encodes them into meaningful embeddings. TCR2vec was jointly pretrained by MLM and the newly proposed SPM via a multi-task pretraining procedure to gain a contextual understanding of TCRs and to enable the preservation of sequence similarity in embedding space. To showcase the effectiveness of TCR2vec, we assessed it in a supervised and an unsupervised task that closely relate to TCR’s recognition ability.

In the supervised classification task, we demonstrated that the embedding generated by TCR2vec largely outperforms other approaches that are based on *k*-mers of TCR or CDR3 sequence with additional V and J gene information. Besides, with only the MLM used in the pretraining phase, the model could not capture the information from sequence similarity that is useful for inferring the binding target ([23, 26]). As a result, it shows lower classification performance. To further manifest the robustness of TCR2vec, we also compared it to state-of-the-art models that either use only the CDR3 or the full-length TCR in predicting the binding of TCR and epitope. We show that TCR2vec surpasses these advanced models, indicating that TCR2vec can partly negate the need for complex prediction models, as a simple SVM and a two-layer MLP can achieve high accuracy.

In the task of clustering TCRs, we validated the utility of TCR2vec in two types of clustering frameworks. For the first type of clustering strategy based on pairwise distances, TCR2vec achieves the best performance with the purity-retention curve lying above such curves of other methods (Fig. 4A). Besides, the improvement over TCRdist indicates that distances in vector space are more informative for clustering TCRs with reduced computation time via vectorization. We also demonstrate that TCR2vec can facilitate GIANA, a representative of the two-step clustering framework (Fig. 4B). The improvements brought by TCR2vec in both classes of clustering frameworks indicate that embeddings produced by TCR2vec are more informative and can be easily grouped into different functional subsets. We hope that TCR2vec could be applied in other existing or ongoing clustering frameworks with enhanced performance.

Note that, we do not consider the alpha chain of TCR in the current study, as most of the previous studies focus on the beta chain ([13, 23, 27]). More importantly, the availability of TCR data with both alpha and beta chain information in existing databases is still limited (e.g., only 18.7% of the TCR-epitope data in our collected dataset contains the information of paired chains). A future update of TCR2vec is expected when more paired data are generated and curated. Besides, the performance of TCR2vec in the classification task could be further improved by applying a more complex classification head (e.g., the Gaussian process model) or combining with advanced CNN-based models since the output tensor **X**_*out*_ is a natural input for CNNs. Also, TCR2vec could be applied to more applications such as inferring the thymic selection pressure ([6]). We leave these possible improvements and explorations in future work.

In conclusion, we present a deep representation learning framework specific for TCR sequences in this study to pave the way for future applications of full-length TCR sequences. We have demonstrated the effectiveness and robustness of TCR2vec in two downstream tasks, and showed that CDR3 only partially encodes the TCR’s function. Therefore, TCR2vec can facilitate the development of more accurate and robust machine learning models, providing new insights for understanding the underlying complex sequence pattern of TCRs.

