## Supplementary Materials for "TCR2vec: a deep representation learning framework of T-cell receptor sequence and function"

### Supplementary Material

#### 1 Text S1

***EigDecom* [1].** In this approach, the embeddings for epitopes are the concatenation of the features of amino acids computed through the eigendecomposition of the BLOSUM62 matrix. First, in order to get the embeddings of amino acids, we performed eigendecomposion to the BLOSUM62 matrix (with gap and normalization) in the form of $B = VSV^T$. The row vectors of $V$, indexed by the amino acids $a$ from the BLOSUM62, is the embedding $\phi(a)$ of amino acid $a$. Thus, for any two amino acids $a$ and $b$, we have $\phi(a)S\phi(b) = [B]_{ab}$. To embed epitope sequences, they were first aligned to the fixed length of 15 by introducing gaps in the middle of the sequence. Then the embedding of a given epitope $x$ was the concatenation of the embeddings of its amino acids: $x = [\phi(a_1), \phi(a_2), ..., \phi(a_{15})]$. We set the size of the embeddings for amino acids to be 8 such that each epitope is transformed into a vector with a size of 120.

***AtchleyAE* [4].** This approach is based on an autoencoder model, which uses convolutional layers for reconstruting the atchley factors of the input amino acid sequences. Each input epitope was first transformed into a 2D map with the shape of $L \times k$, where $L$ is the padded length and $k = 5$, which is the number of atchley factors for each amino acid. The features of the dense layer at the middle of the autoencoder are extracted as the embeddings of epitopes. We followed the same training settings of its original publication and retrained it on 362,456 unique epitope sequences collected from Mei *et al.* [5] to learn the sequence pattern of epitopes. Also, the size of the middle layer was set to 120.

#### 2 Text S2

**KMER** [7]. In this approach, the input sequence is first broken into $k$-mers. The amino acid sequence is considered as a sentence and the $k$-mers as words. Then a doc2vec model [2] model is applied to learn the $k$-mer pattern by predicting the $k$-mer from its surrounding context $k$-mers. The KMER model was trained on *Emerson data* with the window size of 2 and $k$ set to 3. We used the trained model to embed full TCR sequence by averaging the embeddings of its non-overlapping $k$-mers.

**DeepTCR** [6]. DeepTCR is a VAE model for learning a joint representation of a TCR by its CDR3 sequence and V/D/J gene usage. The genes are represented by one-hot encodings. The middle of the VAE is extracted as the representation of the input TCR (in the form of CDR3 + V + J). We trained DeepTCR on *Emerson data* using the default training parameters.

#### 3 Text S3

In the finetuning process, the parameters in TCR2vec will get updated via the backpropagation. Thus, TCR2vec can better adjust its parameters to capture the fine-grained information of the specific task. For finetuning on the TCR-epitope prediction task, we again used a two-layer MLP taking the concatenation of the embeddings from TCR2vec and *EigDecom* as input and then make predictions. We finetuned TCR2vec for 20 epochs with a cosine annealing schedule [3] for adapting the learning rate.

#### 4 Text S4

To avoid remembering the training set for prediction and perform a more robust model evaluation, we filter the test set based on the Levenshtein similarity score: for a pair $(t_i, e_i)$ in the test set, if $t_i$ possess a levenshtein similaity score $L_{sim}(t_i, t_i^{'})$ with any TCR $t_i^{'}$ in the training set, then this pair will be filtered out. Note that the Levenshtein distance is computed for the

CDR3 sequence since the full TCRs possess small pairwise levenshtein distances (only small proportion of TCR is within variable domain). The levenshtein similarity score is defined as:

$$L_{sim}(t_1, t_2) = 1 - \frac{Dis(t_1, t_2)}{max\{len(t_1), len(t_2)\}}, \quad (1)$$

where $Dis(t_1, t_2)$ computes the Levenshtein distance (edit distance) between sequences $t_1$ and $t_2$.

### 5 Text S5

Basically, we first aligned and annotated full TCRs according to IMGT numbering schema. Then we replaced each amino acid of a given TCR in the TCR-epitope dataset by alanne (glycine) and used the trained MLP classifier to score the permuted sequence. Then we substracted the permuted scores at each position to the original predicted score and then took the absolute value of the score difference. These difference values were divided by the original scores and were defined as the relative score change (%). We performed this scanning procedure for all TCR-epitope pairs and averaged the relative score change at each residue to obtain the final results shown in Fig. 5D.

### 6 Supplementary Figures

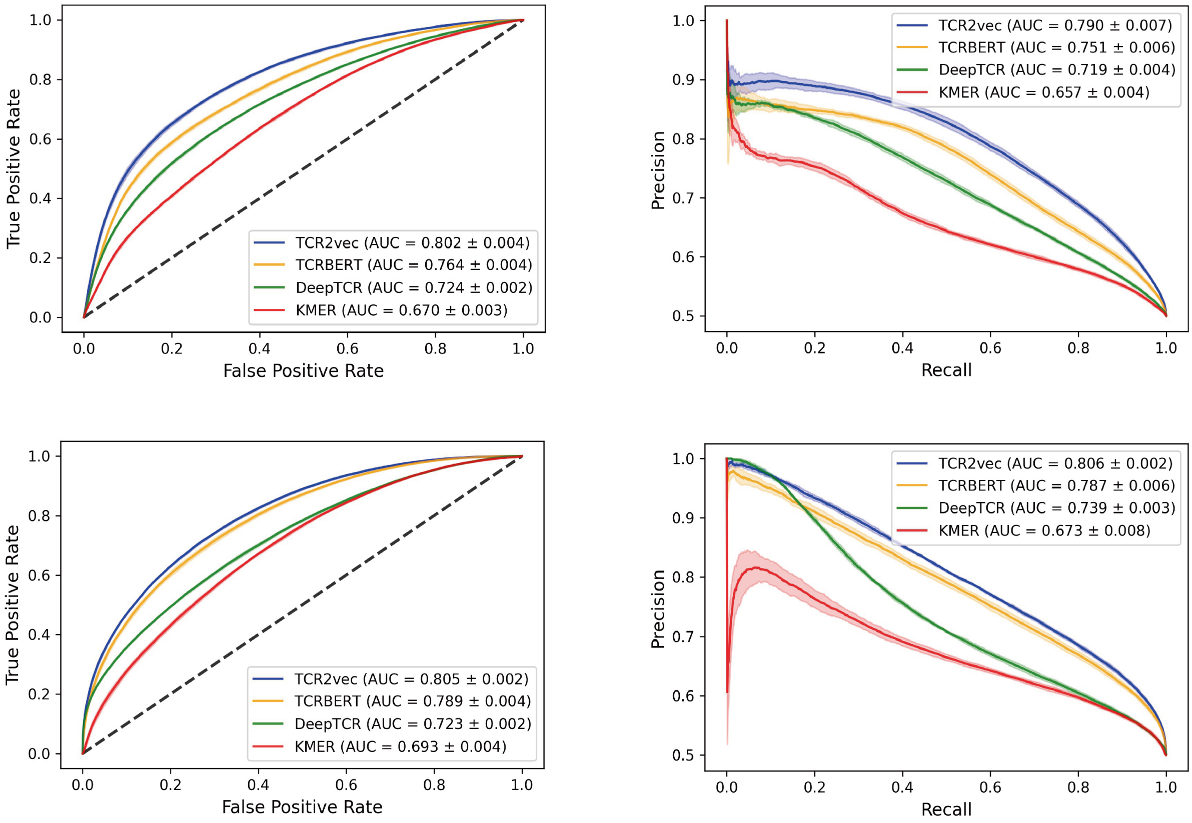


Figure S1: The prediction performance of TCR2vec and the comparative embedding methods in the classification task using *AtchleyAE* to embed epitope sequences. The results with the SVM classification head are shown in the top panel; the results with the MLP classification head are demonstrated in the bottom panel.

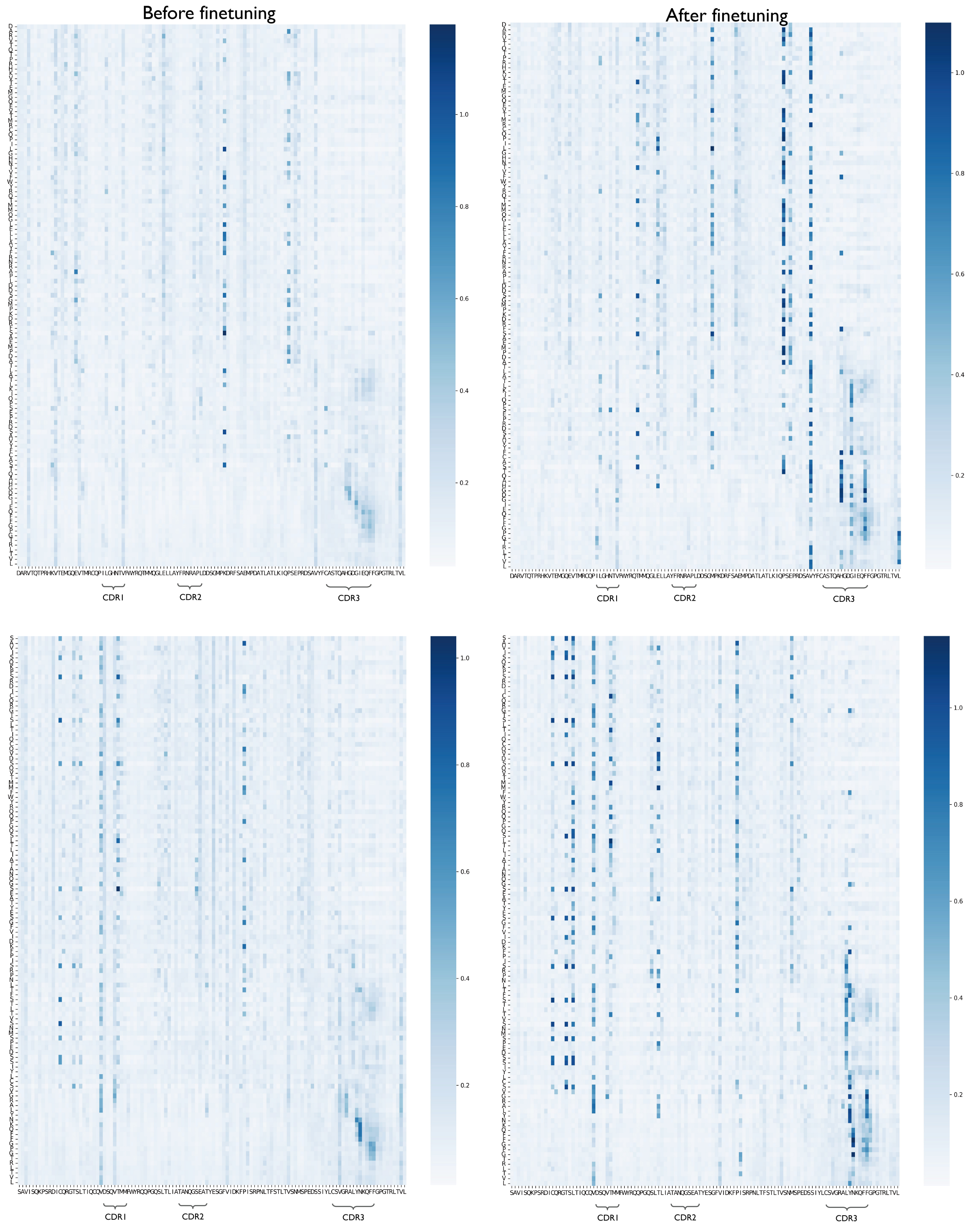


Figure S2: Two additional examples of the general attention map before and after finetuning TCR2vec on the classification task. The results from attention maps further strengthen the conclusion that only limited information is encoded in the CDR3 region.

[3] Ilya Loshchilov and Frank Hutter. Sgdr: Stochastic gradient descent with warm restarts. *arXiv preprint arXiv:1608.03983*, 2016.

[4] Tianshi Lu, Ze Zhang, James Zhu, Yunguan Wang, Peixin Jiang, Xue Xiao, Chantale Bernatchez, John V Heymach, Don L Gibbons, Jun Wang, et al. Deep learning-based prediction of the t cell receptor–antigen binding specificity. *Nature Machine Intelligence*, 3(10):864–875, 2021.

[5] Shutao Mei, Fuyi Li, Dongxu Xiang, Rochelle Ayala, Pouya Faridi, Geoffrey I Webb, Patricia T Illing, Jamie Rossjohn, Tatsuya Akutsu, Nathan P Croft, et al. Anthem: a user customised tool for fast and accurate prediction of binding between peptides and hla class i molecules. *Briefings in Bioinformatics*, 22(5):bbaa415, 2021.

[6] John-William Sidhom, H Benjamin Larman, Drew M Pardoll, and Alexander S Baras. Deeptcr is a deep learning framework for revealing sequence concepts within t-cell repertoires. *Nature communications*, 12(1):1605, 2021.

[7] Kevin K Yang, Zachary Wu, Claire N Bedbrook, and Frances H Arnold. Learned protein embeddings for machine learning. *Bioinformatics*, 34(15):2642–2648, 2018.
